# Assessing the role of fungal diversity in decomposition: A meta-analysis

**DOI:** 10.1101/2021.09.29.462096

**Authors:** Sytske M. Drost, Annemieke van der Wal, Wietse de Boer, Paul. L.E. Bodelier

## Abstract

Fungi play an important role in carbon - and nutrient cycling. It is, however, unclear if diversity of fungi is essential to fulfill this role. With this meta-analysis, we aim to understand the relationship between fungal diversity and decomposition of plant materials (leaf litter and wood) in terrestrial and aquatic environments. The selection criteria for papers were the presence of a fungal diversity gradient and quantification of decomposition as mass loss. In total 40 papers met the selection criteria. We hypothesized that increase of fungal species will result in stronger decomposition, especially in species poor communities. Both artificial inoculated and naturally assembled fungal communities were included in the analysis in order to assess whether manipulated experiments are representative for field situations. We found a significant positive effect of increased fungal diversity on decomposition. However, in manipulated experiments this relationship was only positive when a control treatment of one fungus was compared with multispecies communities. This relationship became negative when comparisons of higher initial richness (at least two fungal species as “control”) were included. In contrast, under natural field conditions increased fungal diversity coincided with increased decomposition. This suggests that manipulated experiments are not representative for field situations. Possible reasons for this are discussed. Yet, both in manipulated and field experiments, environmental factors can influence diversity – decomposition relationships as indicated by a negative relationship of increasing C:N ratio on the effect of fungal diversity on decomposition. Overall, our results show that fungal diversity can have an important role in decomposition, but that design of experiments (manipulated or field) and quality of the plant material should be taken into account for interpretation of this diversity-functioning relationship.

## Introduction

Understanding the consequences of decreasing biodiversity on the functioning of natural ecosystems is one of the highest research priorities in ecological research (Coleman and Whitman, 2005; Lecerf and Richardson, 2010; Delgado-Baquerizo *et al*., 2017). An important aspect of the biodiversity-functioning relationship is the role of belowground biodiversity on soil functioning such as carbon- and nutrient cycling (Wagg *et al*., 2014; Delgado-Baquerizo *et al*., 2016). Saprotrophic fungi are an important group of soil microorganisms involved in decomposition of organic materials and mineral nutrient cycling (van der Wal *et al*., 2013). It is estimated that 1.5 million fungal species occur worldwide (Hyde *et al*., 2007). They are abundantly present in (undisturbed) ecosystems like grasslands and forest floors, but also in aquatic systems like streams that receive input of terrestrial organic matter (Grossart *et al*., 2019). Due to their hyphal growth form and ability to produce a wide range of polymer hydrolyzing and -oxidizing enzymes, saprotrophic fungi have a key role in the degradation of solid, lignocellulose-rich organic materials (van der Wal *et al*., 2013). The number of fungal species with a predominant saprotrophic lifestyle is tremendous and there are strong differences in their abilities to degrade organic compounds (van der Wal *et al*., 2013). Yet, the importance of this high taxonomical and functional diversity of saprotrophic fungi for decomposition processes is not well understood.

Diversity-functioning relationships are mostly studied in experimental settings after the inoculation of a limited number of fungal isolates on sterile plant residues. Many of these studies have been executed in aquatic experimental settings with terrestrial leaf litter that under field conditions naturally falls into streams (Duarte *et al*., 2006; Pascoal *et al*., 2010; Andrade, Pascoal and Cássio, 2016). Manipulated diversity studies in terrestrial ecosystems are less common and do often involve woody materials (Toljander *et al*., 2006; Wagg *et al*., 2014; Hiscox *et al*., 2016). The diversity gradients range from 1 to a maximum of 16 species, which is a common amount of cultivated fungal species retrieved from aquatic ecosystems (Duarte *et al*., 2010). However, in terrestrial ecosystems much higher diversity levels are found (Deacon *et al*., 2006) and with sequencing techniques even more species are detected, but their function and activity are not yet known (Grossart *et al*., 2019).

Interactions between fungi are an important aspect to be included in the prediction of the effect of increasing diversity on ecosystem functioning. Fungi can compete for the same resources leading to competitive exclusion. This can lead to a reduction of the initial inoculated diversity levels at the end of experiments (Toljander *et al*., 2006). In wood logs, competitive interactions are visible between wood-rot fungi (zones between fungal species as described by van der Wal *et al*. (2013)). When interactions are neutral, fungal species co-occur in the same environment without exhibiting harmful or beneficial effects. Decomposition rates are not expected to be different from the average decomposition of each fungus in monoculture (additive effect). Complementarity and facilitation can lead to increased decomposition. For example, Tiunov and Scheu (2005) showed a positive effect by combining cellulolytic fungi and sugar fungi. The production of cellulase supported the consumption of the released sugars by sugar fungi.

Another aspect that can influence the diversity-decomposition relationship is the chemical composition of the material that is decomposed. Fungal species are adapted to the specific composition of the organic materials they decompose. For example, to overcome lignin barriers in complex organic substrates, degradation (white rot fungi) or modification (brown rot fungi) strategies have been evolved in wood decomposition (Mester, Varela and Tien, 2004). Nutrient availability, lignin content and toxic elements can influence wood and litter decomposition as well as the success and outcome of interactions of fungal species during decomposition.

In general, even though interaction effects can have different directions, it is assumed that diversity is important for decomposition (Baerlocher, 2005; Gessner *et al*., 2010; Hättenschwiler, Fromin and Barantal, 2011). Yet, there is still a debate on the extent of diversity importance as redundancy effects within the community may occur, since many species are able to break down organic matter. We performed a meta-analysis to better understand the relationship between fungal diversity and decomposition. We screened manipulated diversity studies from both aquatic and terrestrial ecosystems including leaf and woody materials. The selection criteria for papers were 1) the presence of an initial fungal diversity gradient and 2) quantification of decomposition as mass loss. Decomposition measured as CO_2_ emissions were not included as fungal interactions can affect carbon substrate use efficiencies of individual fungal species leading to difficulties with interpretation of the relationship between respiration and decomposition (Hiscox *et al*., 2015). The following hypotheses were tested:

1. Diversity increases decomposition as increasing species diversity will lead to increased potential of the community to degrade diverse material (niche differentiation/complementarity).
2. The diversity effect will be flattened off with increasing species richness (redundancy effect with increasing species richness).
3. The diversity effect is expected to be higher in litter or woody materials with lower C:N ratio as substrates with a higher C:N ratio are more difficult to degrade and will require specialized fungi. These fungi have to invest a lot of energy in their specialization (e.g. lignolytic enzymes) and prevent competition from other microbes by creating unfavorable growth conditions (Boddy and Hiscox, 2016), resulting in decreased fungal diversity in substrates with a high C:N ratio.
4. The diversity effect will be reduced when measuring mass loss at later time points as compared to earlier time points within the same substrate; facilitation/complementarity is expected to have larger influence during early stages of decomposition as nutrient content is more diverse leading to a possible increased importance of niche differentiation. Environmental studies that tested decomposition under field conditions were selected as well to understand if the diversity-functioning relationship was different under field conditions with spontaneous developed fungal diversity compared to manipulated fungal diversity in laboratory settings. In these studies, differences in diversity of fungal species are based on differences in natural assembly processes on similar organic starting materials. We therefore additionally hypothesize that:
5. Fungal diversity effects will be larger in field settings compared to experimental settings especially in fungal rich communities. In field settings, natural colonization processes (no artificial inoculation) and changing environmental conditions will result in an increase in possible niches.

## Material and methods

### Literature search

The two main criteria for selection of papers on fungal diversity – decomposition relationships were: (1) decomposition is based on mass loss as % loss of plant material and (2) fungal diversity (richness) differences have been compared (at least 2 levels of fungal diversity per study). Web of Science was used as database for literature search using the following search words: “fungal diversity” AND (“litter decomposition” OR “wood decomposition”). This yielded 142 papers in Web of Science. Papers were included until the 1^st^ of March 2021. Reference lists of the papers were also checked to include articles that were missed by the literature search in Web of Science. We included laboratory experiments with manipulated fungal diversity levels and environmental experiments where the environment created differences in diversity due to differences in colonization processes in replicates or treatments. Studies that were excluded from the analysis compared/examined: (1) contrasting environments or plant residues, (2) decomposition compared between successional stages of the experiment (no diversity differences between treatments but over time), (3) toxicity effects of metals or other harmful compounds. Based on these criteria, we kept 16 manipulated studies and 24 environmental studies. The list of included studies is shown in Table 1.

**Table 1:**
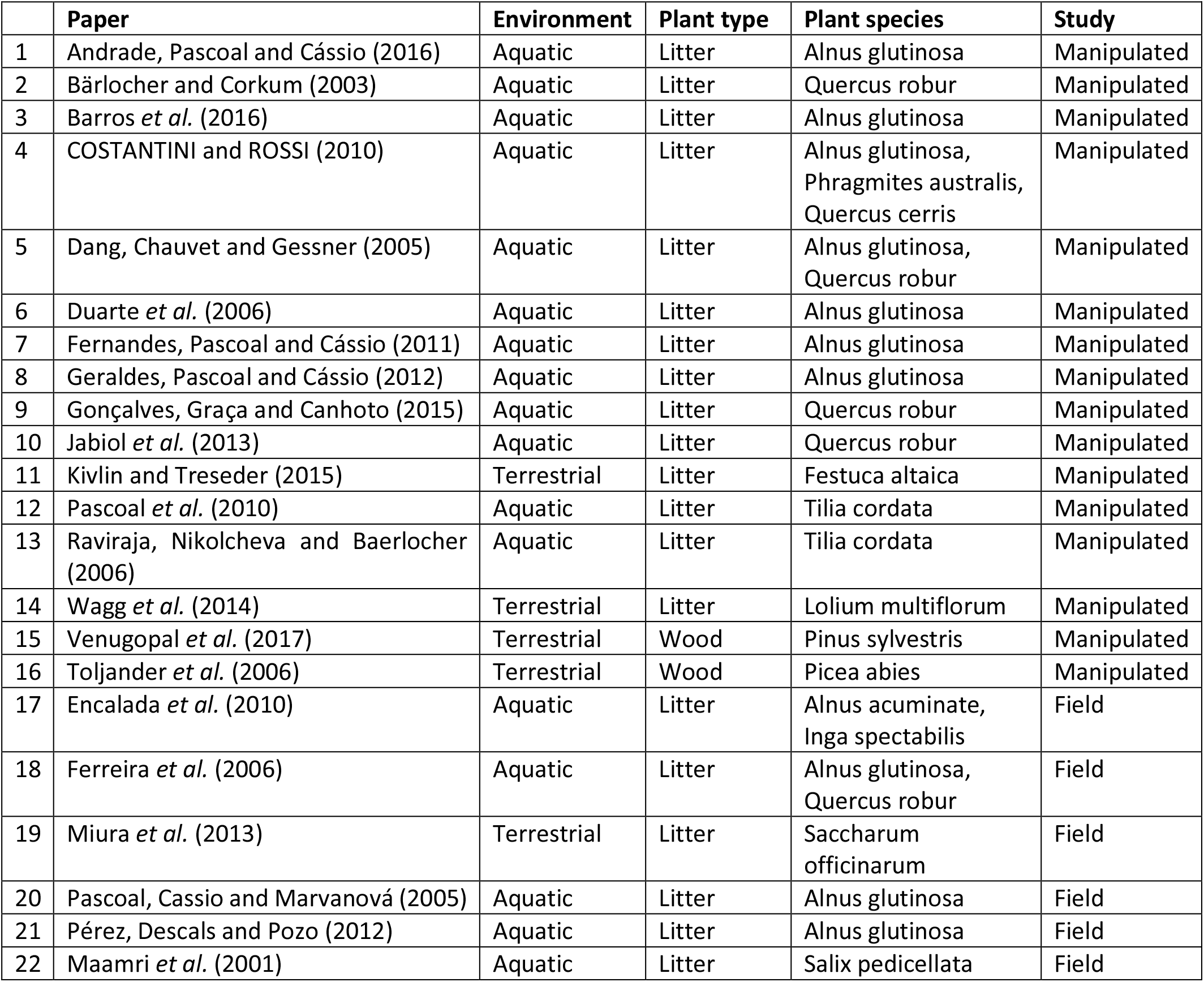
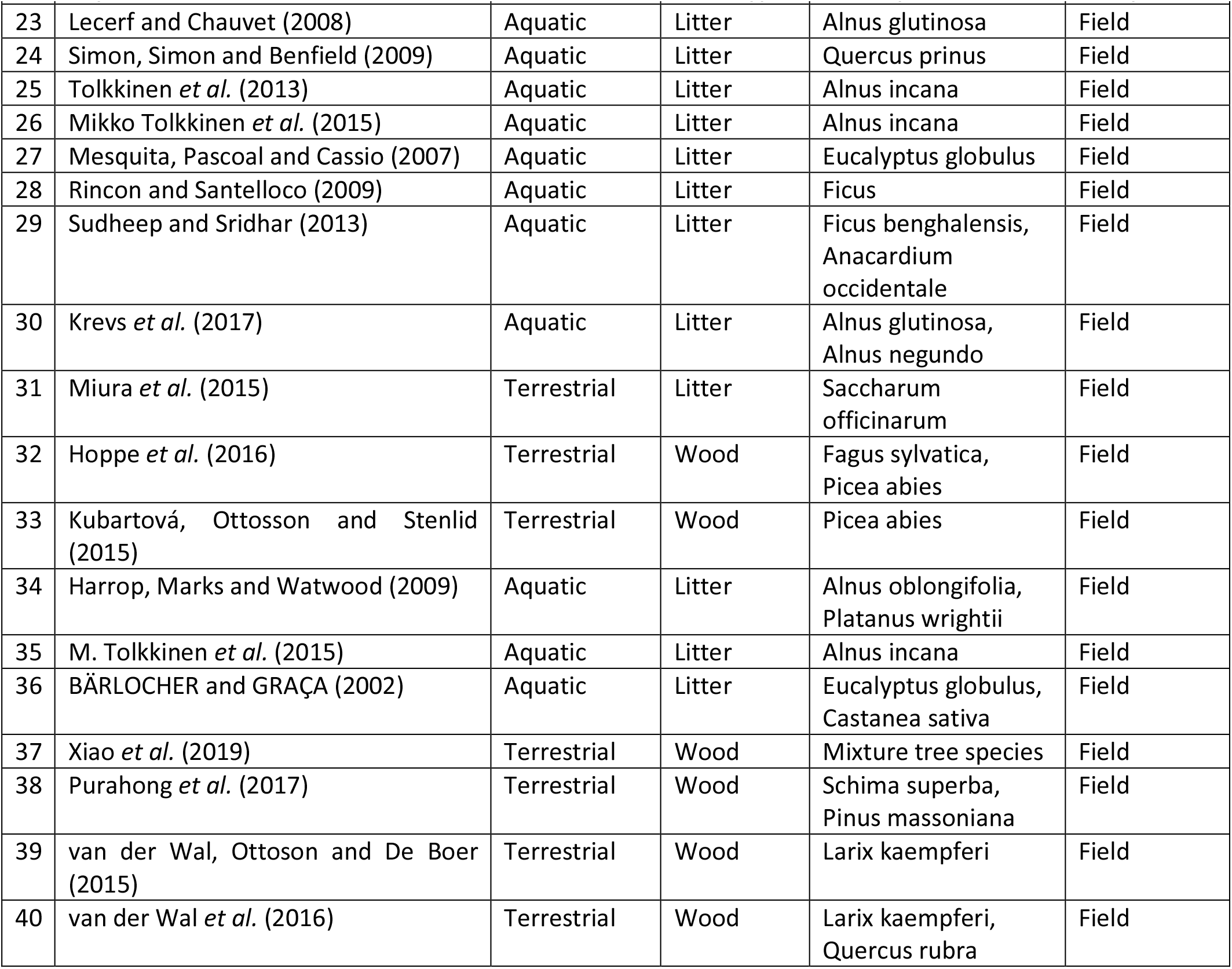
List of papers used in meta-analysis

|  | Paper | Environment | Plant type | Plant species | Study |
| --- | --- | --- | --- | --- | --- |
| 1 | Andrade, Pascoal and Cássio (2016) | Aquatic | Litter | Alnus glutinosa | Manipulated |
| 2 | Bärlocher and Corkum (2003) | Aquatic | Litter | Quercus robur | Manipulated |
| 3 | Barros *et al.* (2016) | Aquatic | Litter | Alnus glutinosa | Manipulated |
| 4 | COSTANTINI and ROSSI (2010) | Aquatic | Litter | Alnus glutinosa, Phragmites australis, Quercus cerris | Manipulated |
| 5 | Dang, Chauvet and Gessner (2005) | Aquatic | Litter | Alnus glutinosa, Quercus robur | Manipulated |
| 6 | Duarte *et al.* (2006) | Aquatic | Litter | Alnus glutinosa | Manipulated |
| 7 | Fernandes, Pascoal and Cássio (2011) | Aquatic | Litter | Alnus glutinosa | Manipulated |
| 8 | Geraldes, Pascoal and Cássio (2012) | Aquatic | Litter | Alnus glutinosa | Manipulated |
| 9 | Gonçalves, Graça and Canhoto (2015) | Aquatic | Litter | Quercus robur | Manipulated |
| 10 | Jabiol *et al.* (2013) | Aquatic | Litter | Quercus robur | Manipulated |
| 11 | Kivlin and Treseder (2015) | Terrestrial | Litter | Festuca altaica | Manipulated |
| 12 | Pascoal *et al.* (2010) | Aquatic | Litter | Tilia cordata | Manipulated |
| 13 | Raviraja, Nikolcheva and Baerlocher (2006) | Aquatic | Litter | Tilia cordata | Manipulated |
| 14 | Wagg *et al.* (2014) | Terrestrial | Litter | Lolium multiflorum | Manipulated |
| 15 | Venugopal *et al.* (2017) | Terrestrial | Wood | Pinus sylvestris | Manipulated |
| 16 | Toljander *et al.* (2006) | Terrestrial | Wood | Picea abies | Manipulated |
| 17 | Encalada *et al.* (2010) | Aquatic | Litter | Alnus acuminate, Inga spectabilis | Field |
| 18 | Ferreira *et al.* (2006) | Aquatic | Litter | Alnus glutinosa, Quercus robur | Field |
| 19 | Miura *et al.* (2013) | Terrestrial | Litter | Saccharum officinarum | Field |
| 20 | Pascoal, Cassio and Marvanová (2005) | Aquatic | Litter | Alnus glutinosa | Field |
| 21 | Pérez, Descals and Pozo (2012) | Aquatic | Litter | Alnus glutinosa | Field |
| 22 | Maamri *et al.* (2001) | Aquatic | Litter | Salix pedicellata | Field |
| 23 | Lecerf and Chauvet (2008) | Aquatic | Litter | *Alnus glutinosa* | Field |
| 24 | Simon, Simon and Benfield (2009) | Aquatic | Litter | *Quercus prinus* | Field |
| 25 | Tolkkinen *et al.* (2013) | Aquatic | Litter | *Alnus incana* | Field |
| 26 | Mikko Tolkkinen *et al.* (2015) | Aquatic | Litter | *Alnus incana* | Field |
| 27 | Mesquita, Pascoal and Cassio (2007) | Aquatic | Litter | *Eucalyptus globulus* | Field |
| 28 | Rincon and Santelloco (2009) | Aquatic | Litter | *Ficus* | Field |
| 29 | Sudheep and Sridhar (2013) | Aquatic | Litter | *Ficus benghalensis*,<br>*Anacardium occidentale* | Field |
| 30 | Krevs *et al.* (2017) | Aquatic | Litter | *Alnus glutinosa*,<br>*Alnus negundo* | Field |
| 31 | Miura *et al.* (2015) | Terrestrial | Litter | *Saccharum officinarum* | Field |
| 32 | Hoppe *et al.* (2016) | Terrestrial | Wood | *Fagus sylvatica*,<br>*Picea abies* | Field |
| 33 | Kubartová, Ottosson and Stenlid (2015) | Terrestrial | Wood | *Picea abies* | Field |
| 34 | Harrop, Marks and Watwood (2009) | Aquatic | Litter | *Alnus oblongifolia*,<br>*Platanus wrightii* | Field |
| 35 | M. Tolkkinen *et al.* (2015) | Aquatic | Litter | *Alnus incana* | Field |
| 36 | BÄRLOCHER and GRAÇA (2002) | Aquatic | Litter | *Eucalyptus globulus*,<br>*Castanea sativa* | Field |
| 37 | Xiao *et al.* (2019) | Terrestrial | Wood | Mixture tree species | Field |
| 38 | Purahong *et al.* (2017) | Terrestrial | Wood | *Schima superba*,<br>*Pinus massoniana* | Field |
| 39 | van der Wal, Ottosson and De Boer (2015) | Terrestrial | Wood | *Larix kaempferi* | Field |
| 40 | van der Wal *et al.* (2016) | Terrestrial | Wood | *Larix kaempferi*,<br>*Quercus rubra* | Field |

Data on decomposition and diversity were extracted from the articles using the online tool WebPlotDigitizer. Data related to plant quality and environment were extracted as well. As most studies did not measure the quality of the plant material used in the experiments, the TRY-database (Kattge *et al*., 2020) was used to get an estimation of the C:N ratio of the plant material.

Treatments with similar diversity levels, but different species composition, were not analyzed separately. Such treatments were pooled before the analysis using the formulas: % *mean mass loss* = *average*(% *mass loss*_1_ + % *mass loss*_2_ + ⋯ + % *mass loss*_*n*_) and standard deviation 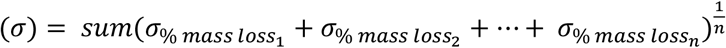 where n is the amount of different treatments with the same diversity level. Some environmental papers measured k decomposition rates and not % mass loss. These values were calculated into % mass loss with the formula: % *mass loss* = 100% − (100% ∗ *e*^−*k*∗*t*^) where k is the decomposition rate and t is the duration of the incubation (in days or years).

### Statistical analyses

Data were analyzed with R (version 4.0.3) with attached packages for analyses and visualization: car, carData, plyr, dplyr, grid, gridExtra, cowplot. The packages metafor and forestplot were used to perform the meta-analysis (Viechtbauer, 2010). Individual effect sizes within each study were estimated by calculating the difference between 2 treatments: *D* = *mass loss* %_*diverse*_ − *mass loss* % _*control*_ and 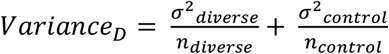 where n is replicates per treatment (diverse or control) (Makowski, Piraux and Brun, 2019). Within each combination, the lowest diversity level was used as “control”. All possible combinations in each study were analyzed leading to multiple effect sizes per study when more than two diversity levels were measured. In total 458 combinations obtained from the 40 selected studies were tested. With a random effect model and the REML method overall effect size was estimated. As most studies had more than one individual effect size (D), we corrected for this within the analysis to prevent overestimation of a single study with more combinations. To determine differences between experimental design (manipulated or field), environment (aquatic or terrestrial) and plant material (litter or wood), these factors were analyzed separately within the analysis. To analyze redundancy effects, an extra factor was added within the manipulated experiments to compare the individual effect size in measured combinations from real control treatments (1 fungal species) and other treatments with a diverse community already (at least 2 fungal species) as lowest diversity level. A forest plot was made to visualize the individual and overall effect sizes and a funnel plot to estimate publication bias (Sterne and Egger, 2001). Regression analysis (with packages ggplot2, ggpubr) was used to assess whether environmental factors and quality of the plant material could explain the magnitude and direction (positive/negative) of individual effect sizes (ES).

Within the environmental studies, seven studies did not analyze treatments, but individual replicates. To estimate the diversity effect on decomposition, linear regression models were used to estimate r and the variance of the relationship between fungal diversity and % mass loss. To be able to compare this approach with the group design of all other studies, the regression results were transformed into d statistics with 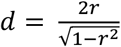 and 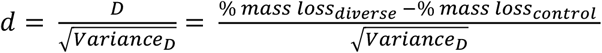 (Nakagawa and Cuthill 2007) where r is the regression coefficient and D is the estimated difference. These estimates of D were added to the other 33 studies to measure the overall effect as described before.

## Results

### Diversity effect

The overall statistical analysis, including all 40 studies, revealed a significant positive effect (D = 1.57±0.41, p < 0.001) of fungal diversity on decomposition (Figure 1), supporting hypothesis 1. The overall effect size was based on individual effect sizes as revealed by a mixed effect model using REML as method (AIC = 3254.2, QE = 14515.9, p<0.001, Table S1). Most of the included studies were performed in aquatic ecosystems and used leaf litter (28 studies). Wood decomposition studies were only done in terrestrial ecosystems (8 studies). Correlation based diversity effects in field studies (7 studies; see M&M) did not change the outcome of the analysis, thus it was not needed to distinguish these field experiments from the other experiments. In the analysis, ecosystem (aquatic or terrestrial), resource (leaf litter or wood), experiment (manipulated or natural assembly) and comparison (control or diverse) were analyzed separately to identify the influence of these experimental factors. The control/diverse grouping was only used in manipulated experiments as in field experiments a fixed control treatment with only 1 fungus was not present. All examined comparisons were statistically significant in explaining differences in the diversity effects on decomposition (QM = 34.72 and p<0.001).

**Figure 1:**
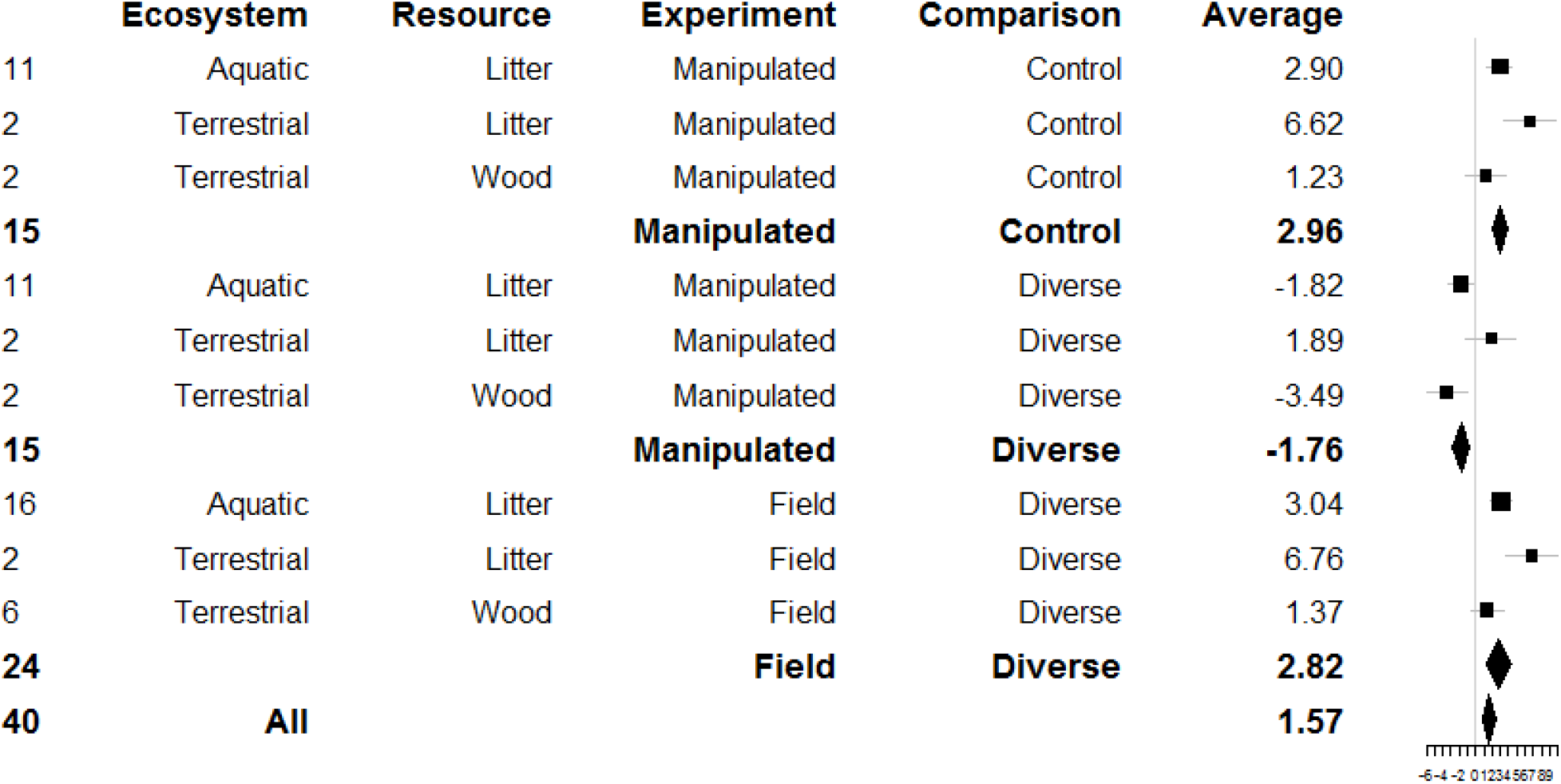
Overall effect size (D) for fungal diversity-decomposition interactions (forest plot), for all studies (All) and different categories within manipulated and field experiments. All data with individual effect sizes of each study calculated as D in forest plot is shown in Figure S1. Numbers in the left column represent the amount of individual studies used in the analysis with a total of 40 papers selected.

In the manipulated experiments, the effect size of increasing diversity was negative (D = -1.76±0.56, p<0.001) when a low diverse community (at least two fungal species) was compared with a higher diversity level, while increasing fungal diversity from one fungus (no diversity) to more fungal species had a positive effect size (D = 2.96±0.46, p<0.001). This indicates a reduced diversity effect with increasing species richness, which is in line with the role of redundancy proposed in hypothesis 2. In field experiments, increasing diversity was positively related to decomposition (D = 2.82±0.78, p<0.001) supporting hypothesis 5. When considering the individual effect sizes (D) of all studies, no significant correlation was seen with the size of the lowest fungal diversity level (control) within each comparison (Figure 2, p = 0.27). This indicates that for field experiments no evidence for a redundancy effect was found.

**Figure 2:**
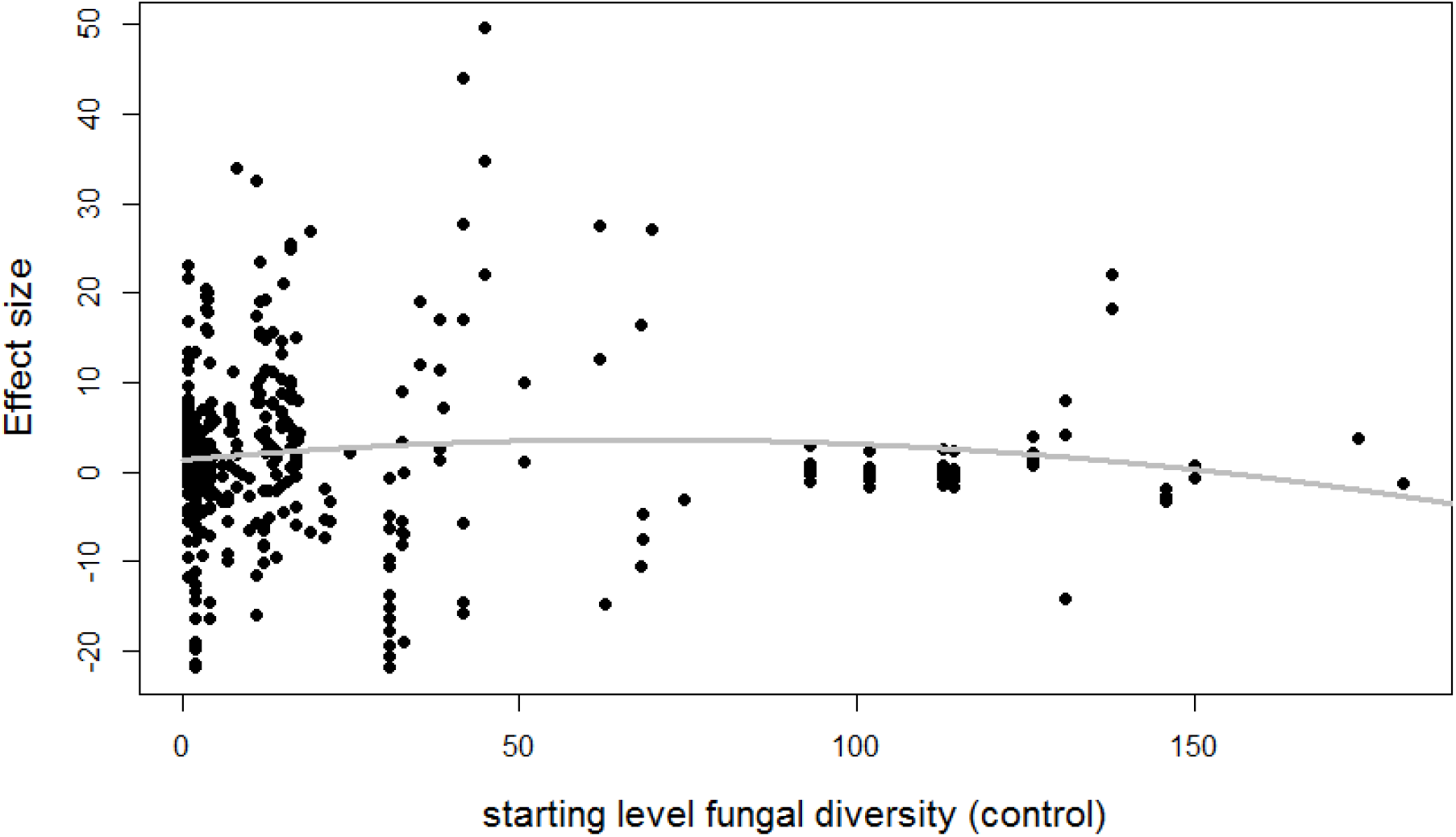
Relationship between effect size (D) and the starting level of fungal diversity within each comparison. Grey line is predicted model based on quadratic relationship (p = 0.27)

Resource type had a significant effect as litter decomposition was significantly increased with increasing fungal diversity (D = 1.65±0.36, p<0.001), whereas this was not the case for decomposition of wood (D = 0.32±1.61, p=0.84). However, the amount of studies with data on wood decomposition was low (8 studies).

To estimate if the selected studies had a publication bias towards studies publishing a significant diversity effect, funnel plot analysis was used to estimate missing studies (Figure 3, Table S2). This resulted in an estimation of missing studies only at the right side of the plot (estimated 110±13.9 studies missing), indicating that in the included studies there was a publication bias for no or negative effects of fungal diversity on decomposition.

**Figure 3:**
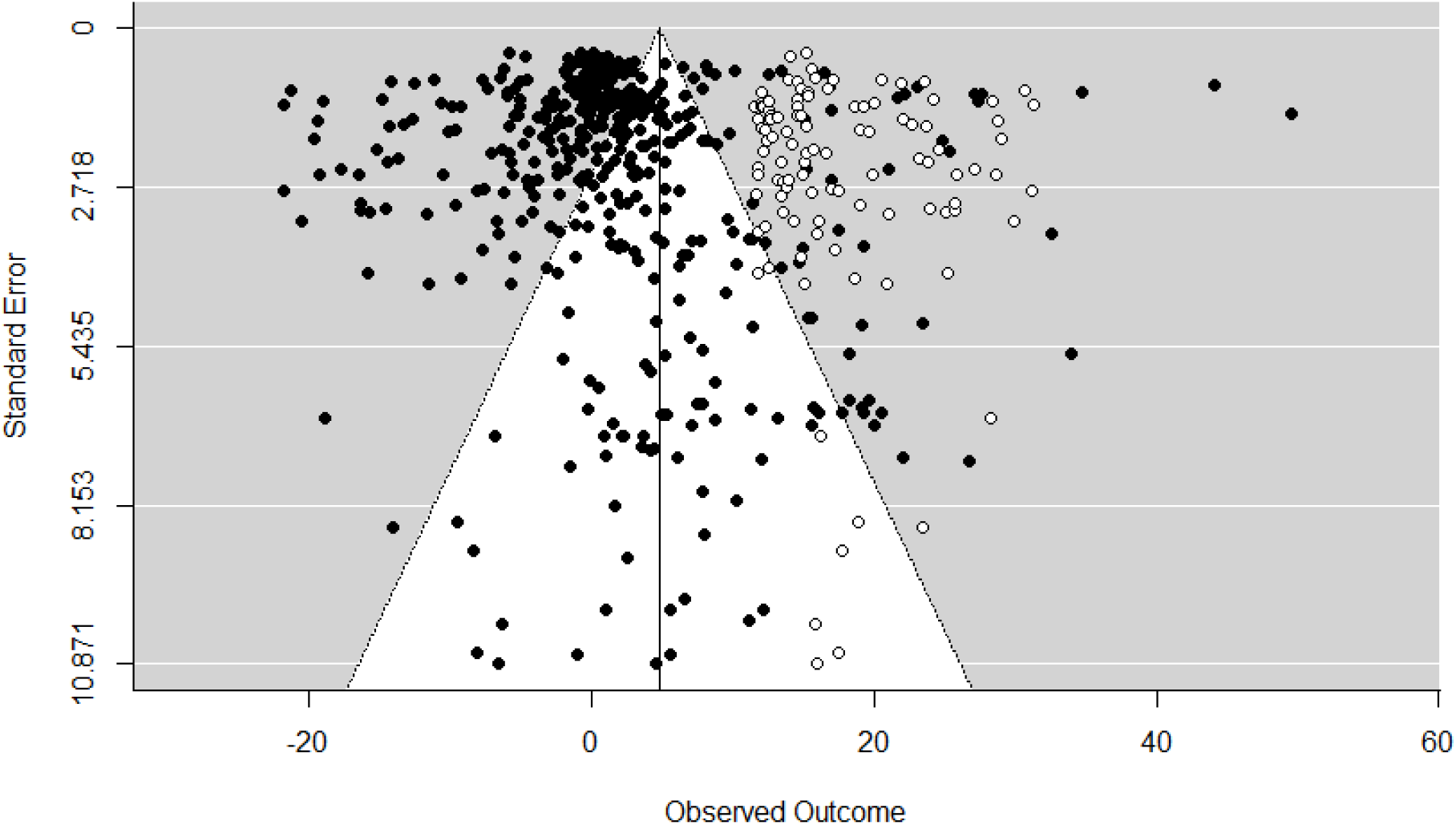
Estimate of publication bias (funnelplot): Individual diversity effect sizes are presented on the x-axis together with the average (black line) and the standard error of the individual effects is presented on the y-axis. Black dots represent individual measurements and white dots indicate possible missing measurements as the distribution of these measurements is expected to be symmetric around the average of all studies.

### Plant quality

Correlation analysis revealed a significant negative relationship between the C:N ratio of plant residues and diversity effect size (r = -0.033±0.016, R = -0.11, p = 0.034, Figure 4), supporting hypothesis 3. The C:N ratio of the plant material was based on an estimate from the TRY-database. For some plant species, however, this data was not available in the database.

**Figure 4:**
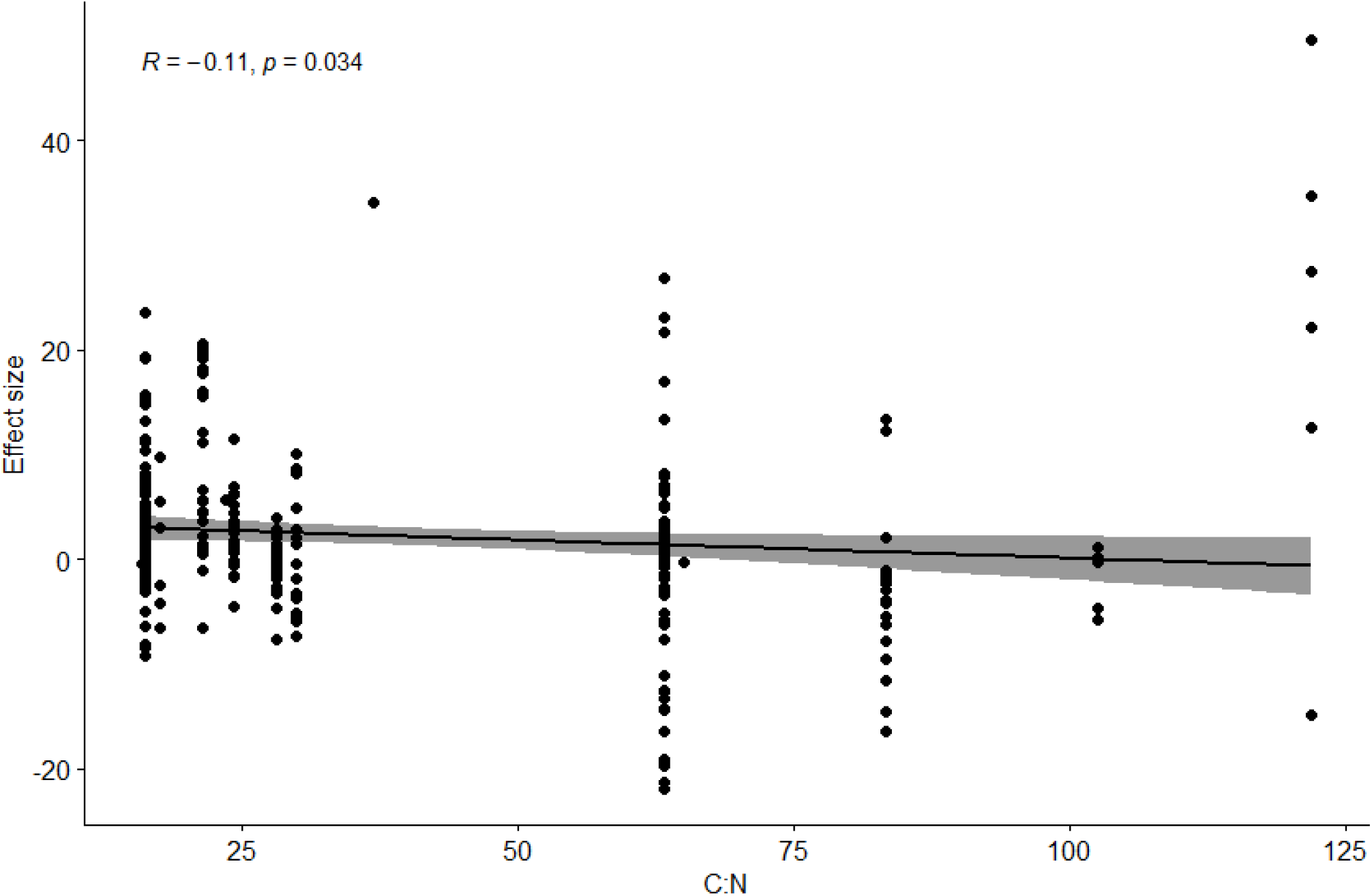
Correlation of estimated C:N ratio of plant material with individual diversity effect size D

### Time effect

To study if time of harvest had an influence on the diversity effect size, the most used plant species was selected to compare different experiments. *Alnus glutinosa* was used in 7 manipulated experiments and 6 field experiments. These experiments were analyzed separately as laboratory conditions are not comparable with field conditions. For both types of experiments the time after which % mass loss was measured did not have a significant negative effect on the individual diversity effect sizes (p = 0.65 and p = 0.98 for manipulated and field experiments respectively, Figure 5), rejecting hypothesis 4.

**Figure 5:**
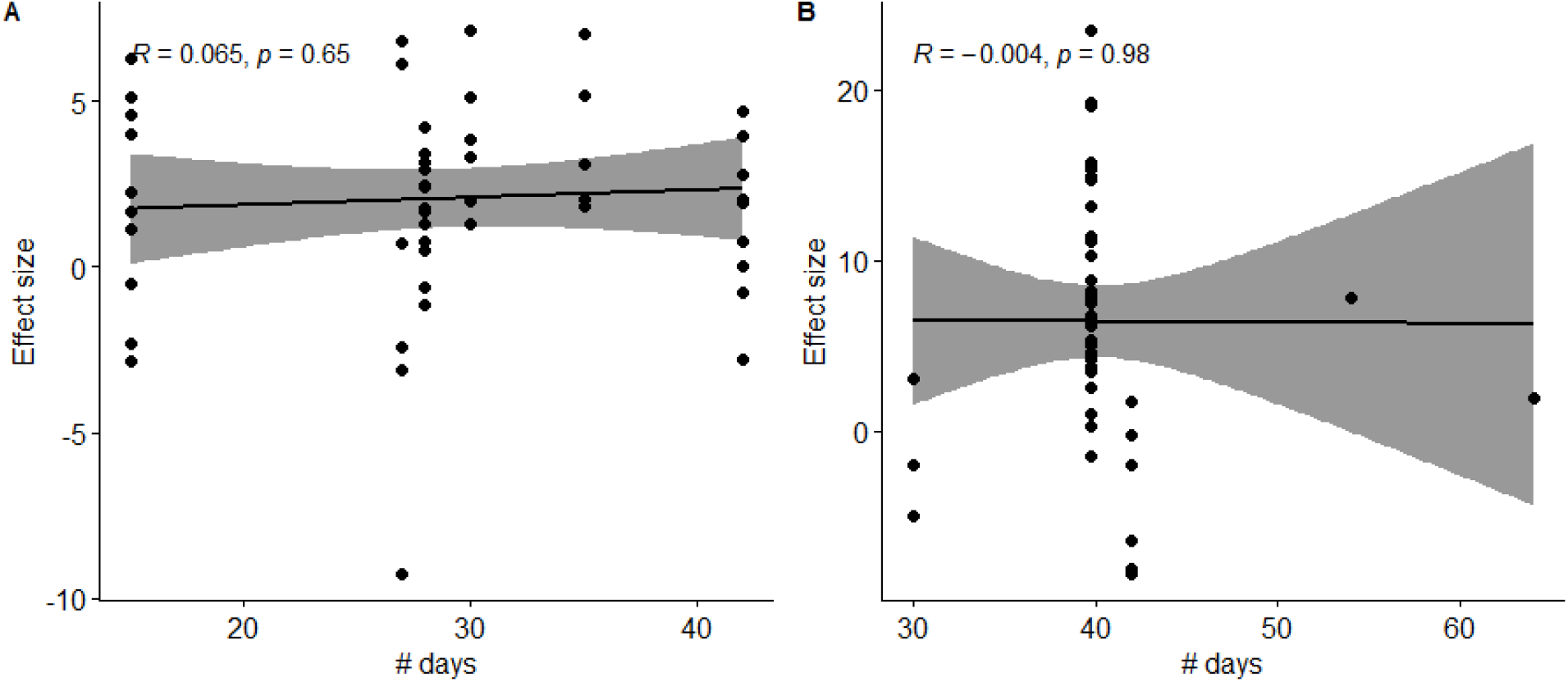
Correlation between time of mass loss measurements and diversity effect size D for 13 studies with leaf litter of *Alnus glutinosa* D. A: manipulated experiments (7) ad B: field experiments (6)

## Discussion

Soil biodiversity plays an important role in ecosystem functioning and nutrient cycling (Thiele-Bruhn *et al*., 2012; Wagg *et al*., 2014; Delgado-Baquerizo *et al*., 2016). As fungi play a prominent role in decomposition and carbon cycling (Creamer *et al*., 2016), the relationship with fungal diversity warrants special attention. Surprisingly, fungal diversity and decomposition are both well studied, but an overall analysis of their relationship has not been addressed. The current meta-analysis examined this both for controlled differences of diversity in experimental settings and for spontaneous assembly differences of diversity in field studies.

In this meta-analysis, we did find an overall positive effect of fungal diversity on decomposition (D = 1.57±0.41, p < 0.001, Figure 1), as stated in hypothesis 1. A positive effect of diversity in manipulated experiments was found when incubations with one fungal species were compared with incubations of two or more fungal species. Comparisons of higher initial richness (at least two fungal species as “control”) with multispecies communities did not show such an effect, supporting the expected redundancy effect (hypothesis 2). However, studies of decomposition of naturally colonized substrates in field sites indicate an overall positive relationship between fungal diversity and decomposition regardless of the lowest diversity levels, thereby rejecting hypothesis 2. These differences between manipulated experiments and natural ecosystems show a possible discrepancy in the representativeness of manipulated experiments for field situations as stated in hypothesis 5. This can be explained in four ways as already discussed by Fukasawa and Matsukura (2021).

First, the fungal diversity levels used to determine the effect of increasing diversity levels on decomposition in experimental settings are the initial levels constructed by the researcher. Competitive exclusion may have leveled off the initial diversity differences in manipulated experiments. For example, Toljander *et al*. (2006) showed that only one or two species existed on decaying wood blocks after 6 months of incubation even though the highest inoculation diversity started with 16 species. Unfortunately, most studies did not take this species loss into account and used the initially inoculated numbers of species to calculate diversity effects on the decomposition as determined at the end of the incubation. This can result in a bias of studies reporting no significant effect of diversity on decomposition as shown in Figure 3. Both competitive interactions leading to a reduction in species and interactions in the remaining species-poor decomposer communities will greatly determine the effect on decomposition (Hiscox *et al*., 2015; Hiscox, O’Leary and Boddy, 2018). In field studies, fungal community composition and decomposition (weight loss) are determined at the same time. Hence, different from artificially manipulated diversity experiments, the relationship between decomposition and fungal diversity in field experiments is not based on the initial, but rather on the final fungal diversity. Therefore, the lack of effect of differences in initial species richness on decomposition in artificially assembled fungal communities cannot be considered as an appropriate test for diversity effects when final richness levels have not been included. Furthermore, in manipulated experiments, a random assembly of fungal species is used as inoculation while in field experiments succession and colonization are not random and can be selected by forces based on functioning, stimulating co-occurrence (Palmer, Stanton and Young, 2003; Dickie *et al*., 2012; van der Wal *et al*., 2013).

Second, the amount of detected species in field experiments is larger than the amount of used species in laboratory conditions (Grossart *et al*., 2019; Fukasawa and Matsukura, 2021). Only one experiment in this meta-analysis used dilution-to-extinction (Wagg *et al*., 2014) as a method to reduce diversity in laboratory conditions. Yet, this method might be more representative for differences in diversity levels in field situations as dilutions are made from naturally assembled fungal communities.

Third, differences in homogeneity and quality of the plant material within manipulated and field experiments can lead to differences in decomposition. In manipulated experiments, the plant material needs to be sterile before fungal communities can be inoculated. These pre-treatments do change the quality of the plant material. Comparisons between partly and fully sterilized material as a method to dilute fungal diversity is therefore not suitable to estimate diversity effects as reported previously (Valentín *et al*., 2014; Muszynski *et al*., 2021). Furthermore, homogenization of the material in laboratory conditions may reduce substrate complexity compared to field conditions. Increasing complexity of the environment and substrate can lead to reduced exclusion during competition and, therefore, lead to co-existence (Lee *et al*., 2019; Chan *et al*., 2021). Even in field situations, reduced resource complexity can lead to reduced diversity of fungal communities (Baessler *et al*., 2014).

Fourth, higher environmental heterogeneity in field experiments as compared to controlled incubation conditions may lead to differences in the relationship between fungal diversity and decomposition. Environmental heterogeneity can lead to increased possibilities for co-existence of fungal species (Bradford *et al*., 2014). Fungal species have specific ranges of abiotic stress tolerance (niche width) (Maynard *et al*., 2019). Therefore, spatial and temporal fluctuations in environmental conditions can facilitate species coexistence which can coincide with increased decomposition (Toljander *et al*., 2006). In this meta-analysis we included both aquatic and terrestrial ecosystems. Even though aquatic and terrestrial ecosystems are different, fungi fulfill an important role as decomposers of terrestrial plant material in both environments. In aquatic ecosystems, an important source of terrestrial plant material consists of fallen leaves and more than 50% of the organic matter in lakes has a terrestrial origin (Wilkinson, Pace and Cole, 2013). Fungal diversity in streams and lakes is simpler compared to terrestrial systems as a possible consequence of reduced environmental fluctuations in temperature and water availability in aquatic ecosystems (Bärlocher and Boddy, 2016; Grossart *et al*., 2019). Even though the presence of these differences, fungal functioning and interactions play similar roles in relationship to the degradation of plant material. However, a limited amount of studies was found that studied decomposition of plant litter in terrestrial ecosystems (4 studies) compared to aquatic ecosystems (28 studies). This bias hampers comparison of both ecosystems. A similar problem accounts for wood decomposition as well. Wood decomposition in aquatic ecosystems is rarely studied even though not only leaf litter but also twigs, branches and even trunks fall into streams as organic matter input (Allan and Castillo, 2007). This limited attention for aquatic wood decomposition is probably due to the assumed very slow decomposition of submerged wood, which has recently been questioned (Ferrer *et al*., 2020). More experimental studies analyzing fungal diversity within decomposing plant material in a diverse set of ecosystems is needed to be able to make better predictions. In addition, by using knowledge from decomposition models that include microbial parameters, it will be possible to better estimate decomposition rates under a changing climate and in different land use types. Until now, only a few models have been developed including microbial traits (Treseder *et al*., 2011; Allison, 2012; Sainte-Marie *et al*., 2021). Our results show that fungal biomass on its own is not enough and fungal diversity should be included in models to better estimate and understand ecosystem functioning. For example, in a modeling approach by Bastida *et al*. (2021) an estimate was made of both microbial diversity and biomass worldwide.

Apart from tested differences in the fungal diversity-decomposition relationship between manipulated and field experiments, we evaluated if plant quality influenced the fungal diversity effect on decomposition. We found a negative relationship between C:N ratio and the effect size supporting hypothesis 3. In the current analysis, it was not possible to quantify the effect of the quality of the plant material in more detail as this information was lacking. Only one out of 40 papers measured C:N ratio. To overcome this problem, we extracted C:N data from the TRY-database (Kattge *et al*., 2020). However, this is an estimation and an average of known measurements, but may not accurately reflect the wood and litter quality used in the experiments. Yet, we did find a negative effect of C:N ratio on the calculated diversity effect sizes (r = -0.033±0.016, R = -0.11, p = 0.034, Figure 4). This suggests that lower plant quality can lead to a reduction of positive effects of fungal diversity on decomposition. Antagonistic forces, such as the production of secondary metabolites and modification of the environment (e.g. reducing pH) to protect occupied space and nutrient sources by the fungal species (Hiscox and Boddy, 2016; Baldrian, 2017), can result in reduced diversity. For example, Baldrian *et al*. (2016) showed that individual wood logs were dominated by one or a few fungal species. Other information about the quality of the plant material, like lignin concentration, can give more insight in the relationship between the plant quality and decomposition (Hall *et al*., 2020). In the selected studies, only a limited amount of data was available for such plant traits. Therefore, other traits related to quality of litter and wood used for the decomposition studies were not included in the analyses.

To test if incubation time had an effect on diversity-decomposition relationships (hypothesis 4), we selected only studies that used *Alnus glutinosa* leaf litter. Within these studies, the differences between harvesting time periods were minimal (ranging from 15 days to 42 days in laboratory conditions and 30 to 64 days in field conditions). This might be the reason for not finding a significant effect and rejecting hypothesis 4. To be able to understand the effect of time on fungal diversity-decomposition relationships, it is better to analyze decomposition at different time points within the same study to account for environmental differences. For example, Fernandes *et al*. (2009) measured decomposition over time, but this study could not be used since the temperature treatments created a diversity effect (exclusion reason 1: contrasting environment or plant material).

In conclusion, fungal diversity is an important parameter to take into account to estimate and understand decomposition. Experimental studies examining fungal diversity effects on ecosystem functioning do not represent natural environments, leading to the need of doing more field experiments to determine fungal diversity-functioning relationships.

## Supporting information

Supplementary table and description of Figure S1

