## Supplementary table and description of Figure S1 for "Assessing the role of fungal diversity in decomposition: A meta-analysis"

*attached as separate figure

Figure S1: All individual effect sizes for each study indicated by the numbers on the left side: first number is publication number, second number is the number of the combination/interaction within each study. Numbers on right side indicate average ± confidence interval. Grey labels indicate average effect size of the individual groups based on the 4 factors (see Figure 1 for average of these groups).

Table S1: Mixed model results (tau^2 = 65.47±4.84, I^2 = 99.85%, H^2 = 670.80, R^2 = 6.59%)

|  | **estimate** | **SE** | **z-score** | **p-value** | **Lower confidence interval** | **Upper confidence interval** |
| --- | --- | --- | --- | --- | --- | --- |
| Intercept | 19.0288 | 3.3442 | 5.6902 | <0.0001 | 12.4744 | 25.5832 |
| Experiment (manipulated) | -4.8658 | 0.9771 | -4.9799 | <0.0001 | -6.7808 | -2.9507 |
| Comparison (control) | -4.7240 | 1.0748 | -4.3953 | <0.0001 | -6.8306 | 2.6175 |
| Ecosystem (aquatic) | 3.7136 | 1.5400 | 2.4114 | 0.0159 | 0.6952 | 6.7319 |
| Resource (litter) | -5.3852 | 1.7876 | -3.0124 | 0.0026 | -8.8889 | -1.8814 |

Table S2: Random effect model, funnel plot analysis with estimate number of missing studies on the right side of 110±13.9 (tau^2 = 99.59±6.40, I^2 = 99.90%, H^2 = 980.61, Q (df = 567) = 30406.67, p<0.001)

|  | **estimate** | **SE** | **z-score** | **p-value** | **Lower confidence interval** | **Upper confidence interval** |
| --- | --- | --- | --- | --- | --- | --- |
| Intercept | 4.6466 | 0.4376 | 10.6193 | <0.0001 | 3.7890 | 5.5042 |
