## Supplementary figures and images for "Assessing the role of fungal diversity in decomposition: A meta-analysis"

### Figure S1

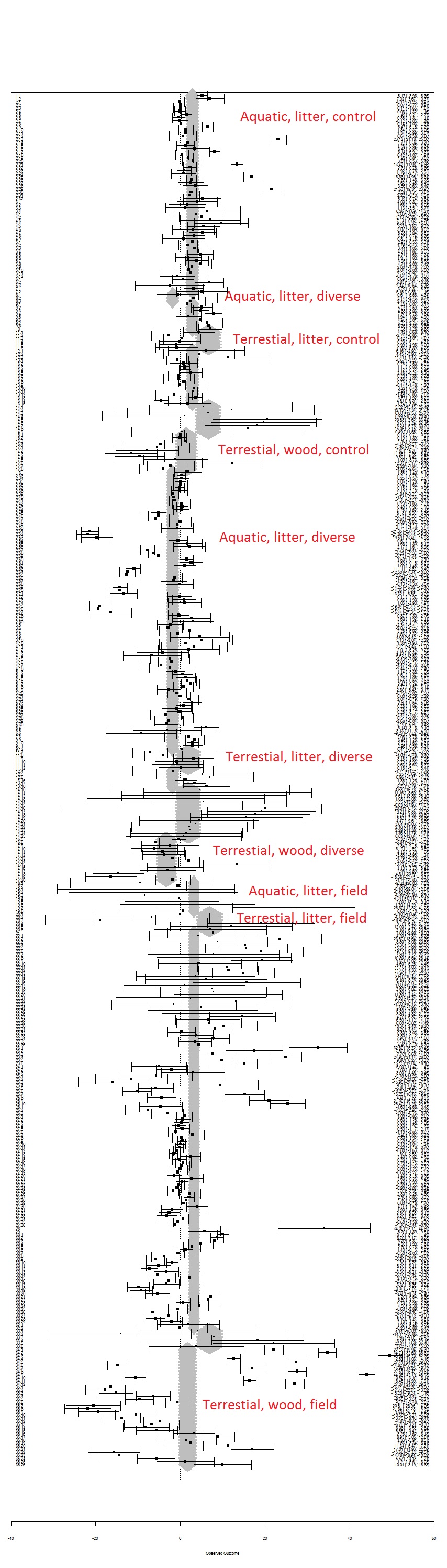
